# Ultra-high field strength electroporation enables efficient DNA transformation and genome editing in nontuberculous mycobacteria

**DOI:** 10.1101/2025.06.04.657955

**Authors:** Daoyan Tang, Minggui Wang, Dan Wang, Danni Yang, Yi Cai, Tao Luo, Jianqing He, Qinglan Wang

**Affiliations:** Department of Respiratory and Critical Care Medicine, West China Hospital, Sichuan University, Chengdu, China; Institute of Respiratory Health, Frontiers Science Center for Disease-related Molecular Network, West China Hospital, Sichuan University, Chengdu, China; State Key Laboratory of Respiratory Health and Multimorbidity, West China Hospital, Sichuan University, Chengdu, China; Guangdong Provincial Key Laboratory of Infection Immunity and Inflammation, Department of Pathogen Biology, Shenzhen University Medical School, Shenzhen, China; Department of Pathogen Biology, West China School of Basic Medical Sciences & Forensic Medicine, Sichuan University, Chengdu, China

## Abstract

Efficient DNA delivery is essential for genetic manipulation of mycobacteria and for dissecting their physiology, pathogenesis, and drug resistance. Although electroporation enables transformation efficiencies exceeding 10^5^ CFU per µg DNA in *Mycobacterium smegmatis* and *Mycobacterium tuberculosis*, it remains highly inefficient in many non-tuberculous mycobacteria (NTM), including *Mycobacterium abscessus*. Here, we discovered that NTM such as *M. abscessus* exhibit exceptional tolerance to ultra-high electric field strengths, and that hypertonic preconditioning partially protects cells from electroporation-induced damage. Using ultra-high electric field strength (3kV/mm) electroporation, we achieved dramatic improvements in plasmid transformation efficiency—up to 106-fold in *M. abscessus*, 83-fold in *Mycobacterium marinum*, and 24-fold in *Mycobacterium kansasii*—compared to standard conditions (1.25 kV/ mm). Transformation efficiency was further influenced by the choice of selectable marker. Ultra-high field strength electroporation also markedly enhanced allelic exchange in *M. abscessus* expressing Che9c RecET recombinases, increasing recovery of gene deletion mutants by over 1,000-fold relative to conventional electroporation. In parallel, oligonucleotide-mediated recombineering for targeted point mutations produced nearly 10,000-fold more mutants under ultra-high field conditions. Together, these findings establish ultra-high field electroporation as a robust, broadly applicable platform for genetic engineering of NTMs. This method substantially enhances transformation efficiency and enables construction of advanced genetic tools—including expression libraries and CRISPRi knockdown libraries—in species that have historically resisted genetic manipulation.

**Importance:** Infections caused by non-tuberculous mycobacteria (NTM), including *Mycobacterium abscessus*, are increasing globally, yet genetic manipulation of these pathogens remains technically challenging due to inefficient DNA delivery and low gene editing success. The ultra-high electric field strength electroporation strategy described here overcomes these barriers, enabling dramatic improvements in both transformation and genome editing efficiency. This advance paves the way for high-throughput functional genomics in NTMs, including the construction of genome-wide knockout, CRISPRi knockdown, and expression libraries. Broad adoption of this approach will accelerate discovery of genetic determinants of virulence and drug resistance, facilitating the development of antimicrobials and vaccines.

---

The genus Mycobacterium comprises nearly 200 species, with nontuberculous mycobacteria (NTMs)—excluding *Mycobacterium tuberculosis* complex and *Mycobacterium leprae*—largely considered environmental and non-pathogenic. However, several NTMs have emerged as opportunistic pathogens, particularly *Mycobacterium abscessus*, which has shown a rising incidence globally(1-3). Its marked intrinsic resistance to antibiotics renders current treatments poorly effective, with cure rates of only 20–50% despite extended multidrug regimens(4). Consequently, *M. abscessus* has been dubbed a “therapeutic nightmare” (5, 6). Genetic manipulation remains central to understanding mycobacterial physiology and drug resistance. While transformation efficiencies in *M. smegmatis* and *M. tuberculosis* via electroporation can exceed 10^5^ CFU/µg DNA, NTMs such as *M. abscessus* (∼10^4^ CFU/µg), *M. marinum* (<10^3^ CFU/µg), and *M. kansasii* (<10 CFU/µg) exhibit significantly lower efficiencies, hindering functional genomic studies including gene knockout and CRISPR interference (CRISPRi) library construction.

In this study, we focused on optimizing electrotransformation efficiency in two NTMs: the rapidly growing *M. abscessus* (ATCC 19977) and the slowly growing *M. marinum* (strain M). Incorporation of glycine during competent cell preparation disrupts peptidoglycan cross-linking, thereby enhancing cell wall permeability (7). Indeed, glycine pretreatment (0.2 M) increased *M. abscessus* transformation efficiency by ∼16-fold for kanamycin-resistant plasmid pQL037 (from 1.5 × 10^3^ to 2.4 × 10^4^ CFU/µg) and ∼7-fold for zeocin-resistant pQL038 (from 5.7 × 10^2^ to 4.0 × 10^3^ CFU/µg; Fig.S1 A-B). Antibiotic selection strongly influenced efficiency, with kanamycin consistently outperforming zeocin. Glycine also improved pQL037 and hygromycin-resistant pQL039 transformation in *M. marinum* by 2-and 8-fold, respectively (Fig.S1D). While previous studies suggested temperature during competent cell preparation affects mycobacterial electrotransformation efficiency—recommending 4°C for fast-growing *M. smegmatis* but room temperature for slow-growing *M. tuberculosis*(7, 8)—we observed slightly higher efficiency at room temperature for both *M. abscessus* and *M. marinum* (Fig.S1A, D). Consequently, subsequent competent cell preparations used glycine pretreatment at room temperature.

We further tested additives known to improve transformation in other organisms(9-12). Addition of 1–3% DMSO to electroporation buffer or 2% to recovery media enhanced transformation of glycine-treated *M. abscessus* and *M. marinum* with pQL037 by 2–3-fold, though no benefit was observed with pQL038 or pQL039 (Fig.S1B, C, E,F). Pretreatment with 0.1 M lithium acetate and 10 mM dithiothreitol— compounds used to increase yeast transformation efficiency—boosted pQL037 transformation in *M. abscessus* ∼4-fold beyond glycine alone, and yielded variable improvements in *M. marinum* (Fig.S1B, C, E, F). Lastly, heat shock at 42°C has been shown to reduce restriction system activity in bacteria, potentially improving electrotransformation(13, 14). Subjecting bacterial cultures to 42°C for 1 hour prior to competent cell preparation improved transformation efficiency of *M. abscessus* 7-fold (pQL037) and 2-fold (pQL038), and of *M. marinum* 6-fold (pQL037) and 8-fold (pQL039) (Fig.S1B, C, E,F). Although DMSO, LiAc/DTT, and 42°C treatments each moderately improved electrotransformation efficiency in both species, none substantially elevated efficiency, nor did they demonstrate significant synergistic effects.

Electrotransformation efficiency typically increases with the applied electric field strength, though elevated fields also raise cell mortality due to membrane damage. In mycobacteria—organisms with notoriously resilient cell walls—standard electroporation is performed at 2.5 kV across a 2 mm cuvette (1.25 kV/mm). This standard field strength (1.25 kV/mm) was employed in our previous experiments. We hypothesized that further increasing the electric field strength beyond 1.25 kV/mm might enhance electrotransformation efficiency in NTMs. To mitigate the expected rise in cell death at ultra-high field strengths, we employed a high-osmolarity electroporation protocol: 0.5 M sorbitol was added to the growth medium, and cells were electroporated in a buffer containing 0.5 M sorbitol, 0.5 M mannitol, and 10% glycerol. These osmoprotectants have been shown to enhance bacterial survival under electrical stress (15). As shown in Fig. 1A-B, the electrotransformation efficiency of *M. abscessus* ATCC 19977 increased progressively with field strength. At 3 kV/mm— the upper voltage limit of the Bio-Rad Micro-pulser—the efficiency for pQL037 and pQL038 rose by 84-and 13-fold, respectively, compared to 1.25 kV/mm. Remarkably, even in the absence of osmoprotective pretreatment, ATCC 19977 tolerated 3 kV/mm, achieving 3.8 × 106 CFU/µg (pQL037) and 7 × 10^4^ CFU/µg (pQL038)—representing 105-and 18-fold increases, respectively. In contrast, the GPL-deficient rough morphotype of *M. abscessus* exhibited increased sensitivity to high field strength. Without osmoprotection, this strain showed only modest gains (5-and 9-fold for pQL037 and pQL038, respectively) (Fig.1E). However, high-osmolarity pretreatment markedly improved outcomes, yielding 20- and 55-fold increases and peak efficiencies of 3.8 × 106 and 6.5 × 10^4^ CFU/µg, respectively (Fig.1E). *M. marinum* also demonstrated robust tolerance to high field strength. At 3 kV/mm, pQL037 and pQL039 transformation efficiencies reached 2.9 × 10^8^ and 1.4 × 10^8^ CFU/µg—83- and 29-fold higher than under standard conditions. Osmoprotective pretreatment provided no additional benefit (Fig.1C-D). Interestingly, *M. smegmatis* mc^2^155 displayed reduced transformation efficiency and increased variability at 3 kV/mm, indicative of greater sensitivity to electrical damage (Fig.1F). Pretreatment with high-osmolarity buffer restored efficiency, leading to 8- and 9-fold increases for pQL037 and pQL039, respectively (Fig.S1G). Notably, this strain lacks several polar lipids—including glycopeptidolipids (GPLs)—present in its wild-type ancestor ATCC 607 (16). Combined with the observation that the GPL-deficient *M. abscessus* strain is also more sensitive to electrical damage, this suggests that an intact mycobacterial cell wall structure is critical for resisting damage induced by ultra-high electric fields. A similar trend was observed in *M. kansasii* SC196, a species with exceptionally low baseline transformation efficiency (64 CFU/µg for pQL037; 2 CFU/µg for pQL038), likely due to plasmid segregation defects linked to the eptC gene (17). At 3 kV/mm, transformation improved to 1.5 × 10^3^ and 38 CFU/µg, respectively (Fig.1G-H). We also assessed whether combining ultra-high electric field strength (3 kV/mm) with DMSO, LiAc/DTT, or heat shock (42 °C) could further enhance transformation efficiency. However, these combinatorial treatments conferred no additional benefit beyond that achieved with high-voltage electroporation alone (Fig.S1H-I)

**Figure 1.**
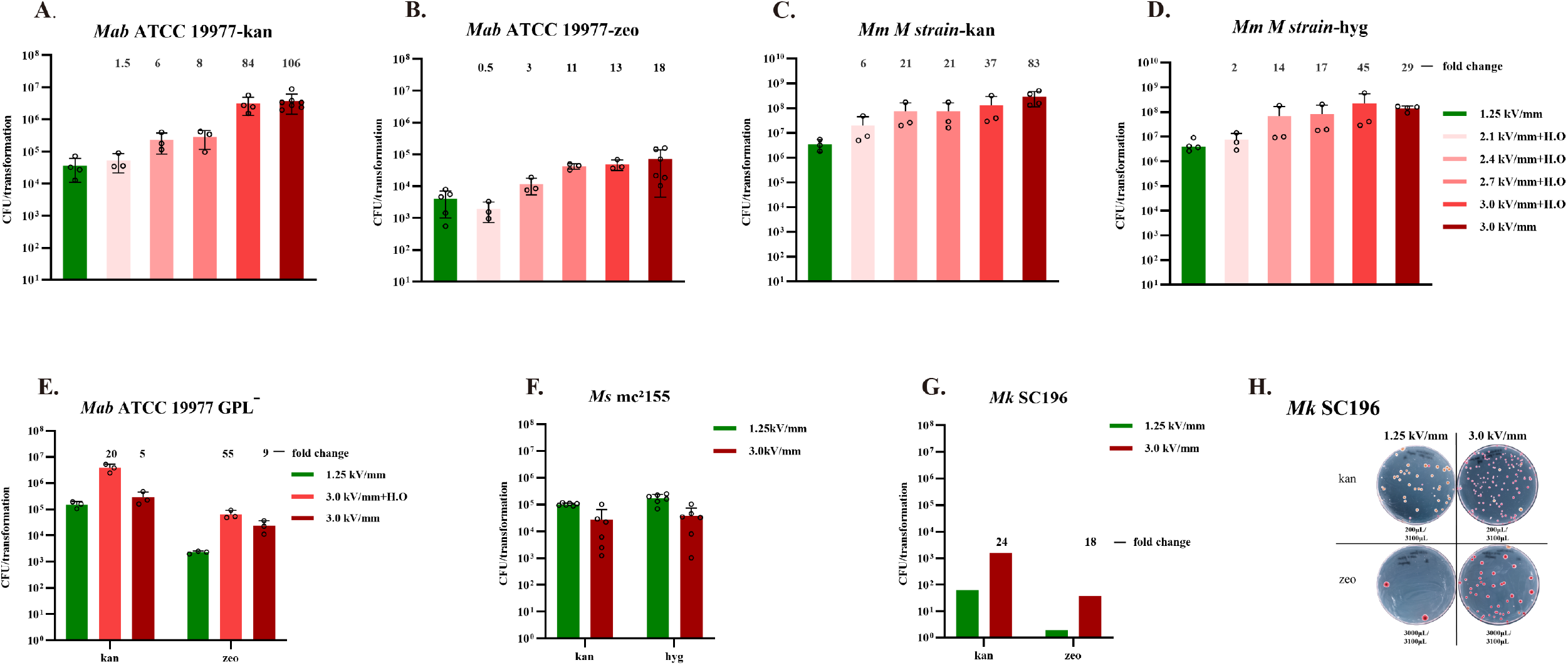
Ultra-high electric field strength electroporation markedly enhances plasmid transformation efficiency in *M. abscessus, M. marinum*, and *M. kansasii*. (A-B) Transformation efficiency of kanamycin-resistant plasmid pQL037 (A) and zeocin-resistant plasmid pQL038 (B) in *M. abscessus* across a range of electric field strengths. Each condition used 100 µl of competent cells and 1 µg of desalted plasmid DNA. Fold increases in colony numbers relative to the conventional field strength (1.25 kV/ mm) are indicated above bars. H.O. denotes hypertonic conditions, achieved by supplementing cultures with 0.5 M sorbitol and adding 0.5 M sorbitol and 0.5 M mannitol to the electroporation buffer in addition to 10% glycerol. (C-D) Transformation efficiency of pQL037 (C) and hygromycin-resistant plasmid pQL039 (D) in *M. marinum* at varying electric field strengths. (E) Effect of ultra-high electric field strength and hypertonic protectants on transformation of pQL037 and pQL038 in a glycopeptidolipid (GPL)-deficient *M. abscessus* strain lacking the mps2 gene. (F) Electroporation efficiency of pQL037 and pQL039 in *M. smegmatis mc*^*2*^*155* under ultra-high electric field conditions. (G) Transformation efficiency of pQL037 and pQL038 in clinical isolate SC196 of *M. kansasii* using ultra-high field electroporation. (H) Representative image of colonies on selective medium following electroporation of *M. kansasii* SC196. Both plasmids express the fluorescent protein mScarlet; purple-red colonies indicate successful transformation. *M. kansasii* electroporation was performed once; all other experiments were independently repeated 3–6 times.

Importantly, by rigorously minimizing salt concentrations in cell suspensions and DNA preparations, we detected no significant arcing events at 3 kV/mm. These results demonstrate that ultra-high field strength electroporation—when combined with appropriate osmoprotective measures in some cases — significantly enhances exogenous DNA transformation efficiency in NTMs such as *M. abscessus, M. marinum*, and *M. kansasii*.

The Che9c RecET recombinase system enables high-efficiency homologous recombination in *M. tuberculosis* and *M. smegmatis*, supporting gene knockouts, promoter replacements, and base editing (18, 19). RecE functions as a 5′→3′ exonuclease, generating 3′ single-stranded DNA overhangs from double-stranded substrates. RecT, a single-stranded DNA-binding protein, promotes strand invasion and homologous pairing. In this system, a plasmid such as pJV53—which expresses recET under acetamide induction—is first electroporated into the host. Allelic exchange substrates (linear DNA with homology arms flanking a selectable marker) are then introduced, with recombinants selected via antibiotic resistance. While RecET recombineering is effective in *M. smegmatis* and *M. tuberculosis*, its application in *M. abscessus* has been limited by extremely low recombination efficiency—only ∼7% of antibiotic-resistant colonies undergo correct allelic exchange (20). Given that our ultra-high field strength (3 kV/mm) protocol enhanced plasmid transformation efficiency in *M. abscessus* by over 100-fold, we tested whether it could similarly improve RecET-mediated genome editing. We targeted six genes (*mmpl4, mmpl6, mmpl7, prcA, uvrB*, and *recD*) for knockout in *M. abscessus* ATCC 19977. To facilitate selection and screening, we modified pJV53 by replacing its kanamycin resistance gene with a zeocin resistance cassette and integrating an mScarlet fluorescent reporter, yielding pJV53-zeo-mScarlet. Allelic exchange substrates (comprising ∼1 kb homology arms flanking a *kanR* gene) were then electroporated (using both ultra-high and standard field strengths) into pJV53-zeo-mScarlet recombinant *M. abscessus* strain after induction of recET expression. Across all six targets, ultra-high field electroporation yielded >10^5^ kanamycin-resistant colonies per transformation, representing 653-to 1,580-fold increases over the standard condition (Fig. 2A-F). In contrast, standard electroporation yielded ∼100 resistant colonies per transformation. To assess recombination fidelity, we randomly selected 23 kanR colonies per gene from each condition for PCR screening. In the ultra-high field group, correct allelic exchange was observed in nearly all colonies (23/23), except for *mmpl6*, which showed a success rate of 96%. By contrast, the standard field group yielded substantially lower efficiencies, with success rates ranging from 13.0% to 43.5% (Fig. 2A-F, Fig. S2A-B). These results demonstrate that increasing the uptake efficiency of linear allelic exchange substrates via ultra-high field strength substantially improves the frequency and precision of genome editing in *M. abscessu*s.

**Figure 2.**
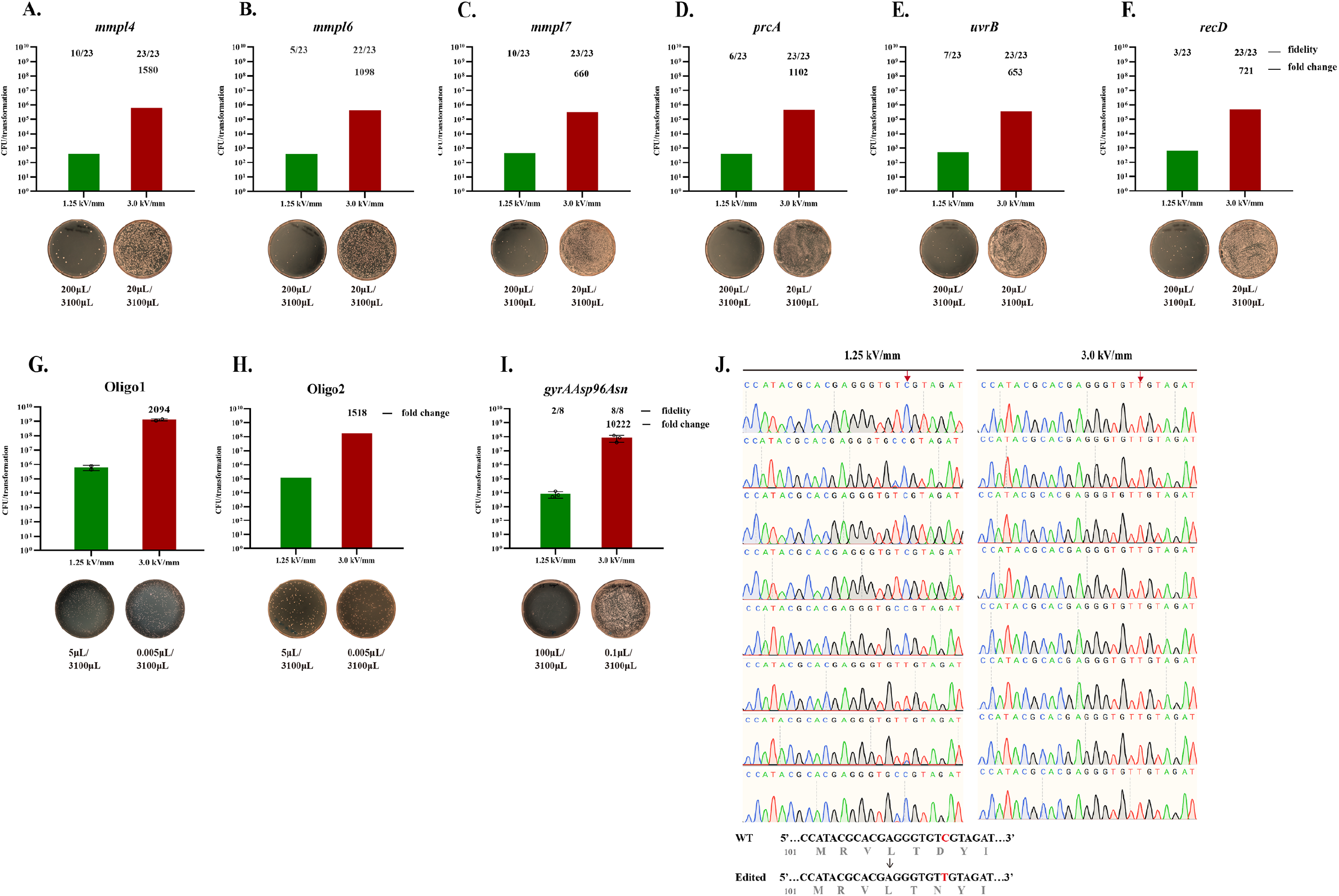
Ultra-high electric field strength electroporation enhances recombination efficiency in *M. abscessus*. (A-F), Gene knockout efficiency is significantly increased using ultra-high electric field strength electroporation of allele exchange substrates in the presence of RecET recombinase. Wild-type *M. abscessus* carrying pJV53-zeo-mScarlet was grown to log phase and induced with 0.2% acetamide to express RecET prior to competent cell preparation. A total of 200 ng of allele exchange substrates targeting six loci—*mmpl4* (A), *mmpl6* (B), *mmpl7* (C), *prcA* (D), *uvrB* (E), and *recD* (F)—were introduced by electroporation under either ultra-high or conventional electric field conditions. Transformants were selected on kanamycin-containing medium. Fold changes in colony numbers (ultra-high vs conventional field strength) are indicated above each bar. For each gene, 23 kanamycin-resistant colonies were randomly selected for verification by colony PCR; successful knockout rates are shown above each bar. Representative selection plates are shown below, with plated culture volumes indicated. (G-H) Oligonucleotide recombineering was assessed using 2 µg of 70-nt oligos (Oligo1, lagging strand; Oligo2, leading strand) targeting a kanamycin repair site in *M. abscessus* harboring the pKM461-dkan-ZeoR plasmid. Electroporation was performed under ultra-high or conventional field strengths, and kanamycin-resistant transformants were enumerated. Fold increases in colony numbers and representative selection plates are shown; plated culture volumes are indicated. (I) Point mutation recombineering was performed using 2 µg of a 70-nt oligonucleotide (Oligo-gyrAAsp96Asn) introducing a c.288C>T substitution in the *gyrA* gene. Electroporation into *M. abscessus* expressing RecT was followed by selection on ofloxacin (80 µg /ml). Fold enrichment and representative plates are shown. (J) Eight colonies from each condition in (I) were analyzed by Sanger sequencing to determine editing accuracy at the *gyrA* locus.

We next evaluated whether this approach could enhance oligonucleotide-mediated recombineering (oligo-recombineering), a strategy that employs RecT to integrate single-stranded oligonucleotides bearing point mutations into the chromosome (18, 21). Oligo-recombineering is efficient in *M. smegmatis* and *M. tuberculosis* (∼1/100 to 1/1,000), but virtually ineffective in *M. abscessus* (<1/106) (18, 21, 22). To test this, we constructed plasmid pKM461-dkan-ZeoR, based on RecT-expressing pKM461 (23), containing a non-functional *kanR* gene disrupted by an E118* nonsense mutation, along with zeocin resistance and mScarlet expression cassettes. Electroporation of a 70-nt oligonucleotide designed to restore *kanR* function yielded markedly more kanamycin-resistant colonies with ultra-high field strength than with standard electroporation. As in previous studies (21), an oligo targeting the lagging strand (Oligo1) produced ∼7-fold more colonies than a leading strand oligo (Oligo2) (Fig. 2G-H). Crucially, ultra-high field strength electroporation, whether using Oligo1 or Oligo2, yielded over 1500-fold more kanamycin-resistant colonies than standard electroporation. To further validate this approach, we targeted a clinically relevant mutation in *gyrA* (c.288C>T; Asp96Asn), known to confer fluoroquinolone resistance (24). Electroporation of Oligo-gyrAAsp96Asn into RecT-expressing *M. abscessus* followed by selection on 80 µg/ ml ofloxacin yielded ∼5 × 10^7^ CFU/2µg oligo with ultra-high field electroporation—over 10,000-fold more than the standard method (Fig. 2I). Sanger sequencing of 8 colonies from each group confirmed the desired mutation in all ultra-high field clones, but only in 2/8 standard-field colonies (Fig. 2J).

In conclusion, efficient delivery of exogenous DNA and genetic manipulation have remained major bottlenecks in NTMs like *M. abscessus*. Here, we show that *M. abscessus* and related species display unexpected tolerance to ultra-high electric field strengths, and that hyperosmotic preconditioning mitigates electroporation-associated damage. By applying electroporation at 3 kV /mm, we achieved a marked enhancement in transformation efficiency in several NTMs, enabling robust gene knockout and oligo-recombineering in *M. abscessu*s. This platform substantially broadens the genetic tractability of NTMs and paves the way for high-throughput approaches, including the development of CRISPRi knockdown and plasmid expression libraries.

